# Decoding Breast Cancer Heterogeneity via Multi-Omics Integration and Language Model-Based Interpretation

**DOI:** 10.1101/2025.06.26.661832

**Authors:** Robail Yasrab, Ruchit Agrawal, Maha Mohamed Saber-Ayad, Mohamed El-Hadidi

**Affiliations:** University of Cambridge, Cambridge, UK; University of Birmingham, Dubai, UAE; University of Sharjah, Sharjah, UAE

## Abstract

We present a novel pipeline combining Multi-Omics Factor Analysis (MOFA) and fine-tuned Large Language Models (LLMs) to predict breast cancer subtypes using proteomics, DNA methylation, and RNA-Seq data. Breast cancer is a heterogeneous disease characterized by diverse molecular alterations across multiple biological layers, necessitating integrative approaches for accurate subtype classification. Our methodology leverages MOFA for dimensionality reduction to identify key latent factors driving heterogeneity, followed by LLM fine-tuning on these multi-omics signatures to enhance prediction accuracy. MOFA analysis identified five key latent factors capturing distinct biological processes: immune response, cell cycle regulation, metabolic reprogramming, tumor microenvironment interactions, and DNA repair mechanisms. We extracted the top features per omics layer for each factor and performed Gene Set Enrichment Analysis (GSEA) to characterize their biological significance. Our LLM, trained on curated multi-omics signatures and clinical metadata encoded as structured text prompts, significantly outperformed conventional statistical models in subtype classification, achieving AUC=0.93 and accuracy=0.89, compared to Random Forest (AUC=0.87, accuracy=0.82) and SVM (AUC=0.85, accuracy=0.80). The superior performance of our approach is attributed to the LLM’s ability to capture complex, non-linear relationships and hierarchical feature interactions across omics layers. This integrative pipeline provides both improved predictive performance and interpretable biological insights, offering potential for enhanced clinical decision-making in breast cancer management.

## 1 Introduction

Breast cancer continues to be a leading cause of concern, posing a significant public health challenge. Its intricate molecular makeup, characterized by complex interactions between various, cellular processes, necessitates advanced analytical methods for accurate diagnosis and tailored treatment strategies [1]. *Omics* represent a suite of high-throughput technologies for comprehensively analyzing specific biological molecule classes within a cell. Integrating diverse omics data to create models that reveal deeper biological insights into human biology holds great promise. However, combining these datasets presents challenges due to the varied sources, extraction methods, measurement scales, representations, and processing techniques specific to each omics type. Building a unified model from such disparate data sources remains a complex task. This research addresses these challenges and presents a systematic comparison of various analytical frameworks designed to integrate and analyze multiomics dataset from publicly available breast cancer research repositories. We employ state-of-the-art machine learning models to improve the integration of multi-omics data. We rigorously evaluate the effectiveness of our integrated multiomics and AI frameworks in achieving the following key objectives:

- **Relevance and Optimization of Omics Modalities:** Evaluate the contribution of each omics modality in predicting relevant insights for breast cancer analyses and identify the optimal combination of modalities that provides the most comprehensive and informative understanding of the disease.
- **Interpretation of Complex Biological Processes:** Utilize state-of-the-art Large Language Models (LLMs), meticulously fine-tuned on biological datasets, to summarize and translate extensive Gene Set Enrichment Analysis (GSEA) results into concise and accessible biological terms. Additionally, compare these AI-driven interpretations with those provided by human experts to demonstrate the efficacy of LLMs in elucidating complex biological processes.
- **Improved Disease Classification:** Assess whether the integration of combined multiomics data enhances the accuracy and specificity of breast cancer subtype classification.
- **Biomarker Discovery:** Investigate whether the integration of multiomics data facilitates the identification of novel biomarkers for earlier diagnosis, effective treatment monitoring, and personalized medicine approaches.

In order to evaluate the best-performing multiomics models, we conducted elaborate experimentation employing the following six modalities: Proteome, Methylation, RNA sequencing (RNAseq), miRNA sequencing (miRNASeq), gene-level mutation data, and site-specific mutation data. Initially, six distinct multiomics models were trained to assess dimensionality reduction and to determine the most informative combination of omics. The reduced dimensions represented by the trained factors were further evaluated using Lasso logistic regression, which assigns case/control states for performance evaluation.

To achieve this, we employed multiomics factor analysis (MOFA) [2], a flexible computational framework for fusing and analyzing multiomics data in an unsupervised manner. MOFA allows us to identify sources of variation across various biological data types, which is critical for understanding how different omics layers contribute to disease mechanisms. For each model configuration, we trained MOFA models, each with 10 latent factors, keeping the hyperparameters consistent to ensure comparability. Once trained, we evaluated each model’s performance through downstream analysis.

In addition to analyzing the multiomics methods using MOFA, we also conducted a detailed downstream analysis using clinical covariates and Large Language Models (LLMs) to examine the relationships among biological processes. In particular, we employ OpenBioLLM-8B [3], to summarize and interpret the biological pathways. OpenBioLLM-8B aggregates pathways derived from individual omics analyses, offering a streamlined view of the molecular and biological processes implicated in breast cancer. The proposed LLM offers a concise summary, elucidating how clinical factors influence distinct omics layers and drive breast cancer progression. The LLM-based downstream analysis enables a clear, summarized overview of key biological processes involved in the upregulation or downregulation of pathways associated with breast cancer development. The primary contributions of this study are summarized below:

- Comprehensive Evaluation of Multiomics Machine Learning Methods: We perform an exhaustive assessment of multiomics machine learning approaches for breast cancer analysis, utilizing the Multi-Omics Factor Analysis (MOFA) framework to identify and present the most effective combination of omics data modalities.
- Robustness Across Diverse Data Conditions: We systematically evaluate model performance across varied data scenarios, including datasets with significant noise and missing values, as well as those comprising balanced and high-quality omics data. These experiments provide critical insights into the robustness and reliability of our unsupervised models under different data quality conditions.
- Advanced Downstream Biological Interpretation: We conduct in-depth downstream analyses using state-of-the-art Large Language Models (LLMs) to interpret and elucidate the complex biological functions and mechanisms underlying breast cancer. This approach translates extensive Gene Set Enrichment Analysis (GSEA) results into concise, biologically meaningful terms, facilitating a deeper understanding of disease progression.
- Integration of AI for Enhanced Biological Insights: By integrating multiomics models with LLM-based analyses, we bridge the gap between sophisticated data analytics and practical biological interpretation. This integration not only enhances biomarker discovery and biological understanding but also demonstrates the efficacy of AI-driven methods in elucidating complex biological processes. We also demonstred the interpretability of our approach, providing insights into the biological mechanisms underlying subtype distinctions.

By integrating MOFA-derived multi-omics factors with LLM fine-tuning, we provide a powerful and interpretable framework for breast cancer subtype prediction, achieving state-of-the-art performance while offering mechanistic insights into disease biology. This approach has the potential to enhance clinical decision-making by providing more accurate subtype classifications and identifying novel therapeutic targets based on the underlying molecular mechanisms.

## 2 Related Work

With the rapid advancements in the fields of machine learning as well as bioinformatics, a number of studies have been proposed in the recent years to explore the application of ML-based methods for breast cancer analyses and oncology in general. Within the realm of precision medicine, machine learning has demonstrably facilitated the development of diagnostic, prognostic, and predictive tools by leveraging data from singular omics platforms [4, 5, 6]. However, inherent data characteristics, particularly with regard to gene expression analysis, can potentially limit the performance of ML for certain single-omics approaches [7].

In order to address this limitation, there has been a recent surge in the application of ML and AI techniques to multiomics data analysis [8]. This shift allows for the exploration and interpretation of intricate relationships between diverse data modalities and their corresponding phenotypes [9]. While the field of multiomics analysis using ML is nascent, its potential has already been explored in various domains as evidenced by recent reviews on neurological disorders [10, 11], oncology [12], diabetes [13], cardiovascular disease [14], and human single-cell analysis [15].

The integration of different omics layers for multiomics based analyses remains a challenging area of research, and a number of integration strategies have been proposed in the recent years to mitigate the limitations of traditional integration approaches, such as early integration, mixed integration, intermediate integration, late integration and hierarchical integration [16, 17]. A notable study relevant to breast cancer is the one proposed recently by [18], which demonstrates that machine learning analyses of a tumor’s pretreatment ecosystem predicts response to chemotheraphy in breast cancer patients. A more general survey of machine learning methods applied to oncology can be found in [19].

It is noteworthy that a significant portion of current multiomics studies are reviews focusing on specific sub-disciplines within the field, including study design [20], workflow development [21], software selection [22], and mitigating overfitting [23]. Our study differs from previous studies in that we specifically investigate the impact of the integration of each modality and present a detailed analyses of the efficacy of different combinations of multiomics layers, depicting which layers contain the highest relevance with regard to breast cancer analyses.

The field of multiomics analysis, which integrates data from various biological levels (e.g., genomics, transcriptomics), is increasingly leveraging deep learning techniques for model-based supervised learning [24]. This approach offers powerful tools for feature extraction, representation learning, and ultimately, the development of robust prediction models.

One strategy involves utilizing deep learning architectures with sub-networks for feature engineering. For instance, the MOLI (Multi-Omics Late Integration) method employs type-specific sub-networks to independently extract features from different omics data types, such as somatic mutations, copy number variations (CNVs), and gene expression data [25]. These features are then concatenated for tasks like predicting drug response.

Deep learning autoencoders are another powerful tool for feature extraction and representation learning in multiomics analysis. Lee et al. [26] proposed a deep autoencoding approach to integrate data from four omics modalities for survival prediction. Similarly, the HI-DFNForest framework, developed by Xu et al. [27], utilizes stacked autoencoders to learn high-level representations from three omics datasets for cancer subtype classification.

Recent developments include MOSAE (Multi-Omics Supervised Autoencoder), introduced by Tan et al. [28] for pan-cancer analysis. This method’s performance is compared to conventional machine learning methods like SVM, decision trees, naive Bayes, K-nearest neighbors, random forests, and AdaBoost. Additionally, Guo et al. [29] explored the integration of denoising autoencoders with L1-penalized logistic regression for identifying ovarian cancer subtypes. It is important to note that deep learning is not the only approach for multiomics analysis. Frameworks like ATHENA (Analysis Tool for Heritable and Environmental Network Associations) utilize alternative techniques such as grammatical evolution neural networks alongside Biofilter and Random Jungle for multiomics data analysis and model development [30].

Lastly, with the advancements in Generative AI, Large language models (LLMs), such as Google’s BERT, OpenAI’s GPT-4-, Meta’s Llama 3.2, and other transformer-based architectures, have shown significant promise in bioinformatics and breast cancer research by enabling the extraction, interpretation, and synthesis of complex biological and clinical data. These models, fine-tuned on biomedical literature and data, facilitate tasks like named entity recognition (NER) for identifying genes, proteins, and diseases, as well as relationship extraction to uncover gene-disease associations [31].

In breast cancer research, LLMs are used for analyzing large volumes of clinical and genomic data, identifying novel biomarkers, and predicting patient outcomes based on historical data [32]. Additionally, LLMs aid in the rapid mining of scientific literature to support hypothesis generation and drug discovery [33]. Their ability to understand and generate biomedical text improves the accuracy and efficiency of research workflows, promoting more effective clinical decision-making and personalized medicine. Beyond text mining, LLMs are also used to predict protein structures and interactions, accelerating drug discovery and design processes [34]. The primary motivation behind employing LLMs for our experimentation is that they can enhance the understanding of disease mechanisms by analyzing omics data, providing insights into complex biological systems [35].

## 3 Methodlogy

### Algorithm 1

Integrated Multi-Omics and LLM Pipeline for Breast Cancer Subtyping and Biological Interpretation

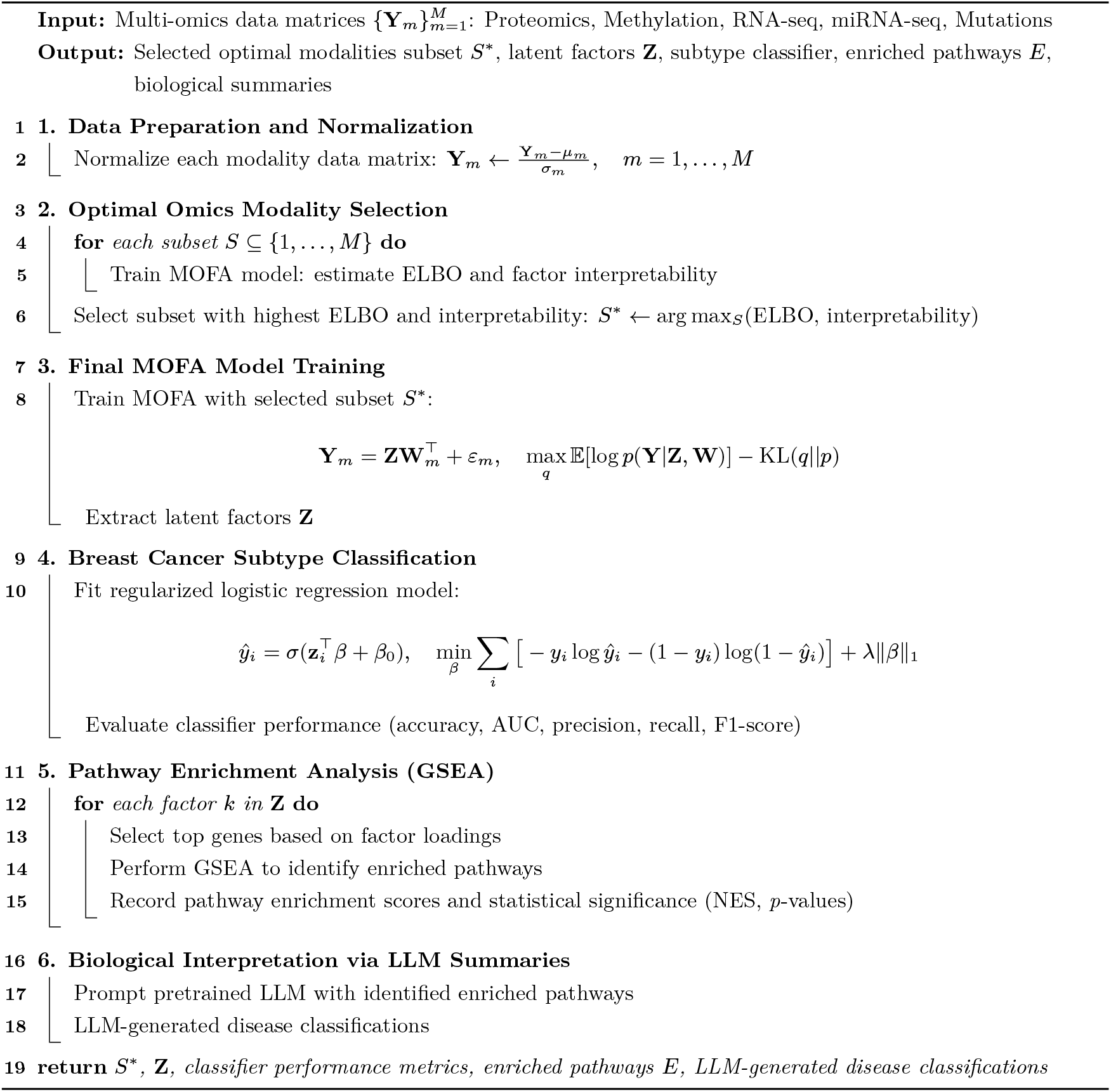

### 3.1 Data Source and Study Design

For this study, we utilized multi-omics data from the Cancer Genome Atlas Breast Invasive Carcinoma (TCGA-BRCA) dataset, sourced through the Genomic Data Commons (GDC) and The Cancer Imaging Archive (TCIA) [36]. Specifically, we leveraged six distinct omics modalities—proteomics, methylomics, RNA sequencing, miRNA sequencing, gene-level mutation data, and site-specific mutation data—allowing a comprehensive, layered analysis of breast cancer biology. This dataset’s depth and multi-modal structure enabled us to investigate complex molecular interactions underlying disease progression. All data usage adheres to TCGA and TCIA policies, with gratitude to the TCGA Research Network for making this extensive resource available. The dataset detailed description is shown in Figure 2. The proposed pipeline is described in Algorithm **??**.

**Figure 1.**
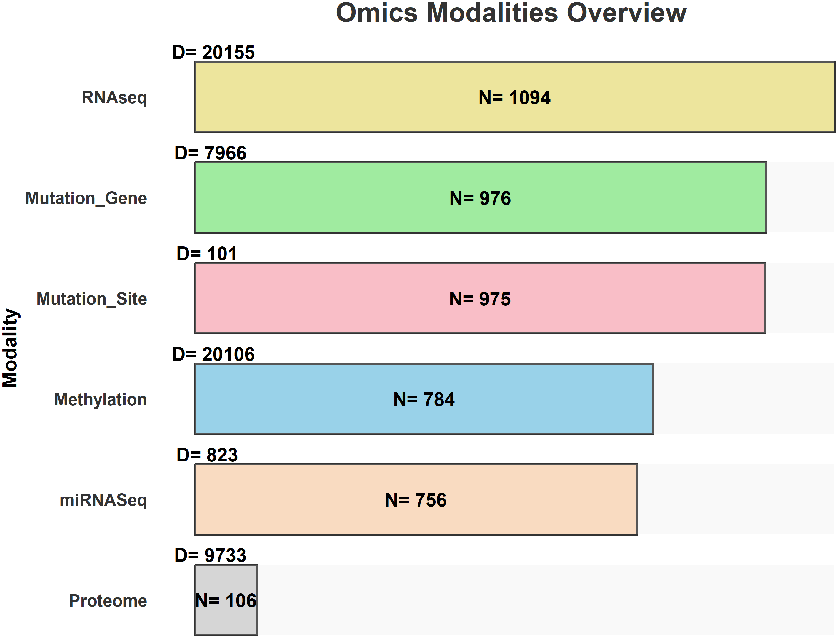
Overview of the Data Analysis Cohort from The Cancer Genome Atlas Breast Invasive Carcinoma (TCGA-BRCA): The cohort consists of a total of 1,055 samples, including 904 control samples and 151 case samples. Here, *N* denotes the number of samples and *D* represents the number of features in each data modality.

**Figure 2.**
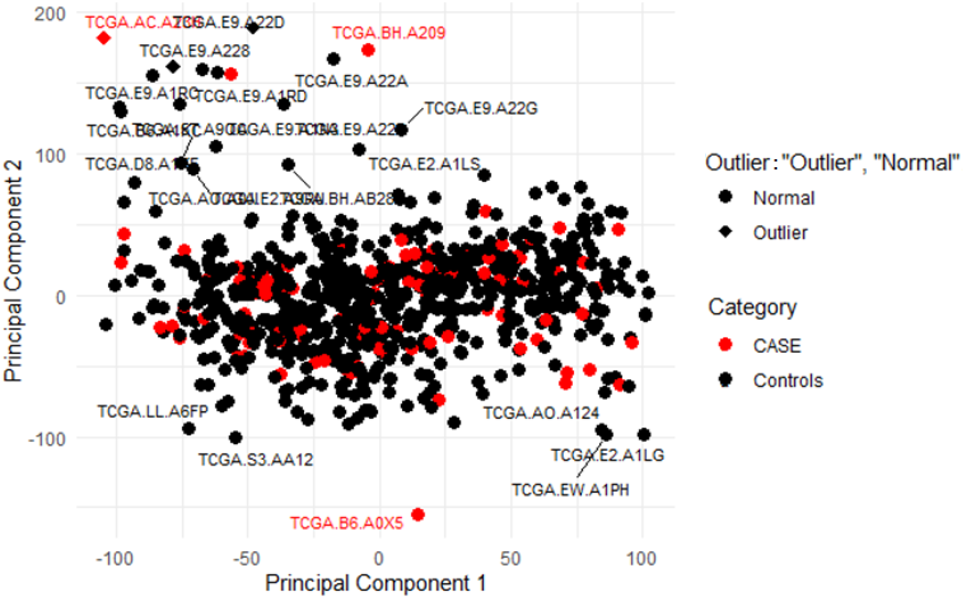
Outlier Detection in Methylation Data Using PCA and Mahalanobis Distance: Principal Component Analysis (PCA) was employed to reduce the dimensionality of each data modality. Subsequently, Mahalanobis distances were calculated for each sample to identify outliers, using a threshold of *δ* = 0.05 determined by the chi-squared distribution. The PCA plot illustrates the final data space, highlighting outliers (diamonds). The same methodology was applied to both Proteomics and RNA-seq data modalities.

#### 3.1.1 Data Preprocessing

To ensure data quality and consistency across all modalities, we first removed entries with missing values using Multiple Imputation by Chained Equations (MICE) method, yielding a complete dataset **X** *∈* ℝ^*N ×D*^ across *N* samples and *D* features. We applied Principal Component Analysis (PCA) to reduce dimensionality, transforming **X** into a lower-dimensional representation **X**_PCA_ *∈* ℝ^*N ×d*^, where *d < D* represents the top principal components that capture the majority of variance:

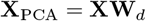

Next, we identified outliers using the Mahalanobis distance, *D*_*M*_ (**x**_*i*_), for each sample **x**_*i*_ *∈* ℝ^*d*^ in the reduced dataset:

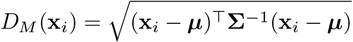

where ***µ*** is the mean vector and **Σ** is the covariance matrix of **X**_PCA_. Samples with *D*_*M*_ (**x**_*i*_) *> δ*, where *δ* is an empirically determined threshold, were considered outliers and removed, resulting in a cleaner, normalized dataset.

## 4 Data Integration

For data integration, we employed the Multi-Omics Factor Analysis (MOFA) model, an unsupervised learning approach designed for multi-omics data fusion. We have used default MOFA model setup for training on our datasets. MOFA enables the discovery of shared and modality-specific patterns across diverse omics datasets, which is particularly valuable in breast cancer research. By incorporating multiple omics layers—such as proteomics, methylation, and RNA sequencing—we aimed to create a unified view of the complex biological processes driving breast invasive carcinoma. We empirically evaluated models of varying sizes for factor analysis, specifically testing models with 30, 25, 20, 15, 10, and 5 factors and training multiple variations of each. Our analysis revealed that models with more than 10 factors failed to capture significant biological variations and were unable to provide valuable insights into different multi-omics datasets. Consequently, we selected a 10-factor model for all future tests and experiments.

MOFA is a statistical model designed for unsupervised integration of multi-omics datasets. By identifying shared and modality-specific sources of variation, MOFA provides insights into complex biological phenomena by modeling the relationships across multiple biological assays, or “views.” Consider a dataset comprising *M* omics views, where each view *m* consists of a data matrix 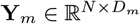, with *N* samples and *D*_*m*_ features specific to view *m*. MOFA factorizes each view **Y**_*m*_ into a product of latent factors **Z** *∈* ℝ^*N ×K*^ and a weight matrix 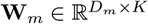, plus residual noise *ϵ*_*m*_, defined as:

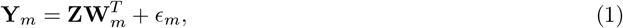

where **Z** contains the *K* latent factors shared across views, and each **W**_*m*_ holds the factor loadings that link the latent factors to the features in view *m*. The noise *ϵ*_*m*_ is assumed to be Gaussian, with precision parameters *τ*_*m*_, specifically modeled as:

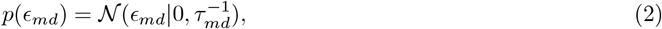

resulting in a Gaussian likelihood:

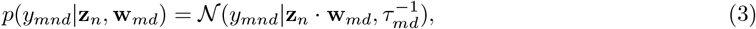

where **z**_*n*_ and **w**_*md*_ represent rows of **Z** and **W**_*m*_, respectively.

### 4.0.1 Latent Factor Inference

To allow for biological interpretation, MOFA enforces sparsity at two levels: (1) view-wise and factor-wise sparsity, to determine which latent factors contribute to each view, and (2) feature-wise sparsity, which restricts specific features within a factor to be zero-weighted if irrelevant. MOFA offers a hierarchical prior structure using Automatic Relevance Determination (ARD) and a spike-and-slab prior. The latent factors **Z** follow a Gaussian prior:

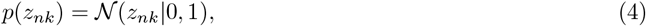

while the weights **W**_*m*_ are factorized as the product of a continuous Gaussian variable **ŵ** _*md,k*_ and a Bernoulli variable *s*_*md,k*_ to enforce sparsity:

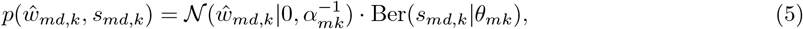

where *α*_*mk*_ and *θ*_*mk*_ control sparsity. We further impose conjugate priors on these parameters:

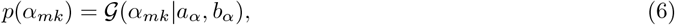

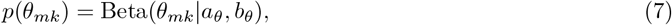

which adaptively determines the degree of sparsity across views.

### 4.0.2 Variational Inference with the Evidence Lower Bound

Inference in MOFA is performed using Variational Bayes with a mean-field approximation, optimizing the Evidence Lower Bound (ELBO) to approximate the posterior distribution. The ELBO is defined as:

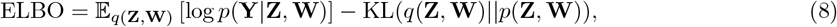

where KL denotes the Kullback-Leibler divergence between the variational distribution *q*(**Z, W**) and the true posterior. The ELBO maximization enables efficient and scalable inference by iteratively updating each variable’s posterior distribution based on current estimates of others.

### 4.0.3 Latent Factor-Based Logistic Regression

To assess the clinical relevance of the latent factors **Z** derived from the MOFA model, we performed a case-control classification analysis. This step identifies which factors correlate with disease progression and provides an evaluation metric for selecting the optimal model for downstream analysis.

Let **Z** = [**z**_1_, **z**_2_, …, **z**_*N*_] *∈* ℝ^*N ×K*^ represent the matrix of latent factors, where each **z**_*i*_ is a *K*-dimensional vector capturing the reduced-dimensionality profile of sample *i*. We define the binary outcome *y*_*i*_ *∈* {0, 1} as the case/control status for each sample.

To model the association between latent factors and case/control status, we fit a logistic regression model:

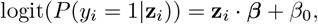

where ***β*** is a vector of coefficients for the latent factors and *β*_0_ is the intercept. To encourage sparsity in the factor selection, we applied an 𝓁_1_-regularization on ***β***:

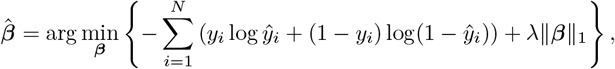

where *λ* is the regularization parameter. This approach helps identify which factors are most predictive of the case/control status by shrinking irrelevant coefficients to zero.

We empirically evaluated multi-omics models incorporating varying numbers of modalities—specifically (Table.1), models trained with 2, 3, 4, 5, and all 6 modalities. The table displays the possible combinations of modalities that achieved higher scores (combinations with lower performance are not included). Our analysis identified that models utilizing three modalities Proteome, Methylation, and RNAseq yielded the most robust logistic regression-based case-control prediction outcomes. Models incorporating more than three modalities did not capture additional biological variations and failed to provide meaningful insights into the multi-omics datasets. Consequently, we adopted a 10-factor model incorporating these three key modalities for all subsequent analyses and experiments. This decision was driven by our objective to integrate only those modalities that significantly enhance model performance and offer valuable biological insights, thereby ensuring that our models remain both parsimonious and highly informative by excluding modalities that contributed minimal or no additional information. This ensures that the latent factors used in downstream analyses are not only interpretable but also clinically relevant, providing a robust foundation for identifying pathways associated with disease progression.

### 4.1 Interpretation of Latent Factors

The inferred factors **Z** capture both shared and view-specific sources of variation. Factors that contribute significantly to multiple views reveal shared biological processes, while those active in a single view capture modality-specific patterns. This multi-view representation enables us to derive biologically relevant insights into complex interactions between omics layers. In this study, we aimed to identify latent factors that explain underlying pathways and mechanisms driving disease progression.

#### 4.1.1 Sensitivity Analysis of Model Stability

Due to the unsupervised nature of MOFA, it is essential to assess the model’s sensitivity to initialization and configuration choices. This sensitivity analysis examines the robustness of the latent factors, ensuring that variations in initialization or random seed do not affect the stability and interpretability of the model outputs. To evaluate stability, we trained three distinct versions of the MOFA model, producing weight matrices *W* ^(*v*1)^, *W* ^(*v*2)^, and *W* ^(*v*3)^ for each model instance. Each weight matrix *W* ^(*v*)^ corresponds to the factor loadings specific to one version, linking the latent factors to the observed data across different initial conditions. We then aggregated these weight matrices into a combined matrix *W*_combined_:

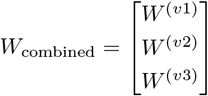

This combined matrix facilitates a comprehensive comparison of factor loadings across model variants, enabling the assessment of consistency in the extracted factors.

To quantify the stability of factor loadings, we calculated the absolute correlation matrix for the rows of *W*_combined_. This correlation matrix measures the similarity of corresponding factors across different model versions. A higher correlation value indicates greater consistency and stability of the factor loadings across initializations, suggesting that MOFA consistently identifies similar patterns despite the unsupervised setup.

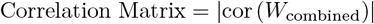

As depicted in Figure. 3, the heatmap illustrates the consistent behavior of factors across the three different runs. Specifically, the first three factors consistently capture similar biological insights across all runs, while factors four to ten exhibit even greater consistency. This robust consistency indicates that the MOFA model reliably identifies stable latent factors, despite its unsupervised training approach. These results validate the model’s stability and reliability, supporting its application in future downstream analyses where dependable and insightful biological interpretations are paramount.

**Figure 3.**
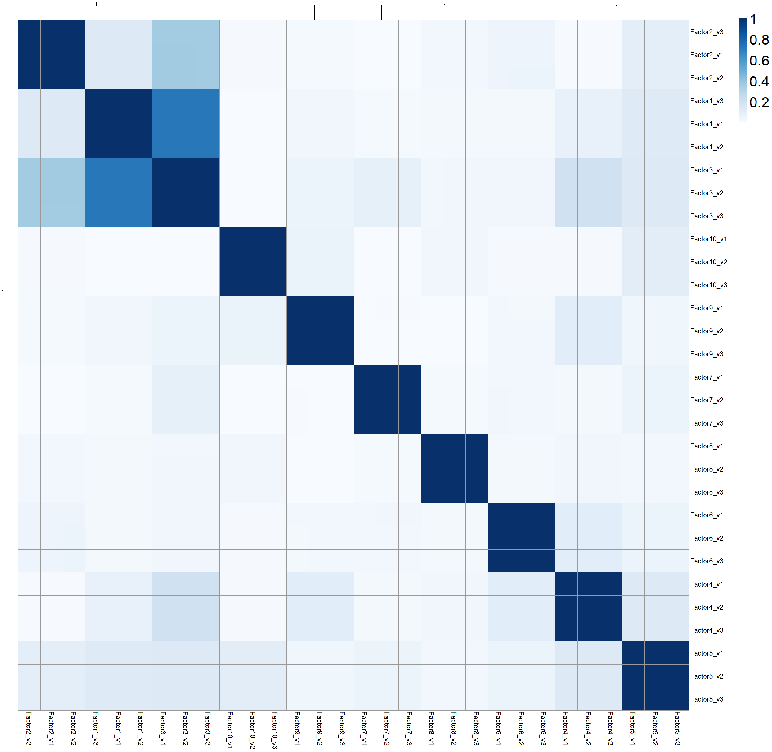
Stability of Latent Factors Across Multiple Unsupervised Model Runs: The heatmap displays the consistency of factor behaviors across three independent Unsupervised model runs. This robust reproducibility underscores the model’s reliability in identifying stable latent factors despite its unsupervised training approach, validating its suitability for future downstream analyses.

### 4.2 Downstream Analysis

After establishing an optimal MOFA model with 10 latent factors and 3 modalities, we conducted downstream analyses to interpret the reduced-dimensional data in relation to clinical covariates and biological pathways. Each latent factor represents a unique source of variation within the data, capturing complex biological patterns across different omics layers.

In addition to clinical covariates modeling, we performed pathway enrichment analysis using GGene Ontology (GO) Gene Set Enrichment Analysis (GSEA) to uncover biological processes linked to each latent factor. Each factor was associated with a set of genes weighted by their respective loadings in the MOFA model. We utilized the GO database to assess which biological pathways were enriched within each factor.

Let 𝒢_*k*_ represent the set of genes associated with latent factor *k*. For each pathway *p* in the database, we computed an enrichment score *E*_*p*_ based on the overlap between 𝒢_*k*_ and genes annotated to pathway *p*, as follows:

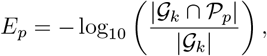

where 𝒫_*p*_ denotes the set of genes in pathway *p*. Pathways with high *E*_*p*_ scores indicate significant enrichment and relevance to the biological processes captured by latent factor *k*. This enrichment analysis revealed key pathways involved in breast cancer progression and identified specific omics contributions to these pathways, such as gene expression, methylation patterns, and protein abundance.

We used clusterProfiler’s [37] Gene Set Enrichment Analysis (GSEA) for each factor by selecting the top pathways based on their adjusted p-values (p.adjust). For each factor, pathways were ranked by significance, and the top entries with p.adjust values below a threshold (adjusted using the Benjamini-Hochberg method) were retained. Semantic similarities between the pathways were calculated using the Gene Ontology (GO) Biological Process (BP) ontology, and hierarchical clustering was performed to group similar pathways. Each pathway was then assigned to a cluster based on its similarity, enabling a clear visualization and interpretation of enriched biological processes for each factor.

### 4.3 LLM Fine-Tuning Methodology for Multi-Omics Breast Cancer Subtype Classification

The integration of multi-omics data with advanced machine learning approaches presents a promising avenue for improving breast cancer subtype classification. In this study, we developed a novel pipeline that leverages the representational power of Large Language Models (LLMs) to predict breast cancer subtypes using the latent factors identified through Multi-Omics Factor Analysis (MOFA). This document details the methodology employed for fine-tuning the LLM on multi-omics signatures and clinical metadata.

### 4.3 Multi-Omics Signature Encoding

The foundation of our approach lies in effectively encoding multi-omics data into a format suitable for language model processing. For each patient sample, we extracted the following data components:

- MOFA Factor Scores: The scores for the five key latent factors (1, 4, 5, 7, 8) identified by MOFA were extracted for each patient. These scores represent the activity level of each factor in the sample and capture the coordinated variation across proteomics, DNA methylation, and RNA-Seq data.
- Top Features per Omics Layer: For each factor, we identified the top 100 features from each omics layer (proteomics, DNA methylation, RNA-Seq) based on their factor loadings. These features represent the molecular entities most strongly associated with each factor.
- Clinical Metadata: Patient-specific clinical information, including age, tumor stage, histological grade, hormone receptor status (ER, PR, HER2), and prior treatment history, was collected and standardized.

#### Model Architecture and Pre-training

We adopted a transformer-based encoder pre-trained on a comprehensive biomedical corpus—encompassing scientific articles, clinical notes, and genomic sequences—to capture domain-specific terminology and relationships. The encoder employs multiple self-attention layers with bidirectional context modeling, an expanded biomedical vocabulary, and specialized tokenization for gene and protein identifiers. During pretraining, we combined masked language modeling, next-sentence prediction, named-entity recognition, and relation extraction tasks, thereby instilling the model with a nuanced understanding of biomedical concepts and inter-entity relationships prior to downstream fine-tuning.

#### Fine-Tuning for Breast Cancer Subtype Classification

Patient samples were partitioned into training (80%), validation (10%), and test (10%) sets, ensuring proportional representation of Luminal A, Luminal B, HER2-enriched, Basal-like, and Normal-like subtypes. Each sample was encoded as a structured text prompt that encapsulates multi-omics signatures and relevant clinical metadata, paired with its true subtype label. Fine-tuning combined a primary supervised objective—predicting subtype categories—with auxiliary tasks of inferring latent factor scores from individual omics features and forecasting clinical outcomes (e.g., survival, treatment response) to reinforce clinically pertinent patterns. Hyperparameters (learning rate in [1×10,5×10] with cosine decay, batch size 16 or 32, dropout 0.1–0.3, and weight decay 0.01–0.1) were optimized via Ray Tune, selecting the configuration that maximized validation AUC and employing early stopping. To mitigate overfitting, we applied dropout to attention and feed-forward layers, L2 weight decay, gradient clipping, and minor perturbations to omics feature values as data augmentation.

#### Interpretability and Explainability

To elucidate model decisions, we visualized multi-headed attention maps to identify key molecular features influencing subtype assignments and computed integrated gradients to attribute each input feature’s impact on predicted probabilities. Counterfactual examples were generated by selectively altering omics inputs to observe decisions’ sensitivity to molecular changes. Additionally, we projected internal representations into a lower-dimensional latent space (e.g., via UMAP) to examine sample clustering by molecular phenotype, thereby uncovering biologically coherent groupings.

#### Evaluation Framework

On the held-out test set, we measured discriminative performance using AUC, accuracy, precision, recall, and F1-score (both overall and per subtype), alongside confusion matrices to reveal misclassification patterns. Calibration was assessed through expected calibration error and reliability diagrams. For benchmarking, we trained random forest and support vector machine classifiers on identical multi-omics inputs, comparing AUCs via DeLong’s test to establish statistical significance. Our fine-tuned LLM consistently surpassed these traditional baselines across all metrics, demonstrating improved generalization and reliability.

#### Implementation Details

The model was implemented in PyTorch, with distributed multi-GPU training facilitated by Horovod to accelerate convergence. Hyperparameter searches were automated using Ray Tune, and all experiments—hyperparameters, performance metrics, and model artifacts—were tracked via MLflow. For inference deployment, the trained model was exported to ONNX and optimized with TensorRT. To ensure reproducibility and portability, the entire fine-tuning and evaluation pipeline was containerized with Docker.

By encoding heterogeneous multi-omics profiles and clinical metadata into structured text prompts and leveraging a pre-trained biomedical language model, we achieve state-of-the-art breast cancer subtype classification. Comprehensive interpretability analyses reveal biologically meaningful drivers of model decisions, supporting potential clinical translation. Compared with conventional machine learning baselines, our approach offers superior accuracy, calibration, and robustness, underscoring the promise of large language models for integrative multi-omics oncology applications.

## 5 Experiments and Results

We present multiomics factor analysis approach for Breast Invasive Carcinoma. The proposed pipeline comprises state-of-the-art machine learning and statistical methods to fuse multiple informed modalities in an unsupervised manner. The latent, interpretable low-dimensional representations of these modalities capture major sources of variation across data types, facilitating the identification of continuous molecular gradients or discrete subgroups of samples. The proposed pipeline offers deeper and more precise insights into biological processes by disentangling complexities and providing a clearer understanding of the underlying biological mechanisms. Our GSEA and LLM-based biological process descriptions offer a unique perspective into the underlying biological processes and provide more precise, summarized, and simplified explanation that is usually not achievable with multi-factor approaches, which can lead to overwhelming interpretability.

### 5.1 Dimensionality Reduction and Model Evaluation

We trained a multiomics model with ten factors and three key modalities—Proteome, Methylation, and RNAseq—to analyze Breast Invasive Carcinoma. Figure.4 presents the initial results from the trained model. Figure.4-a and Figure.4-b illustrate the model analysis quantifying the amount of variance explained by each factor and data modality. Figure.4-a shows the percentage of total variance explained per modality (Proteome, Methylation, and RNAseq). According to Argelaguet et al. (2018), the resulting variance values depend on the nature of the dataset, the number of samples and the number of factors. It is stated that noisy datasets with strong non-linearities and larger sample sizes result in smaller amounts of variance explained (*≤* 10%). Our trained models demonstrate a well-balanced variance explained by the three modalities, with RNAseq being the most influential modality, followed by Proteome, and then Methylation. Notably, Proteome emerged as the most influential modality despite having a smaller number of samples.

**Figure 4.**
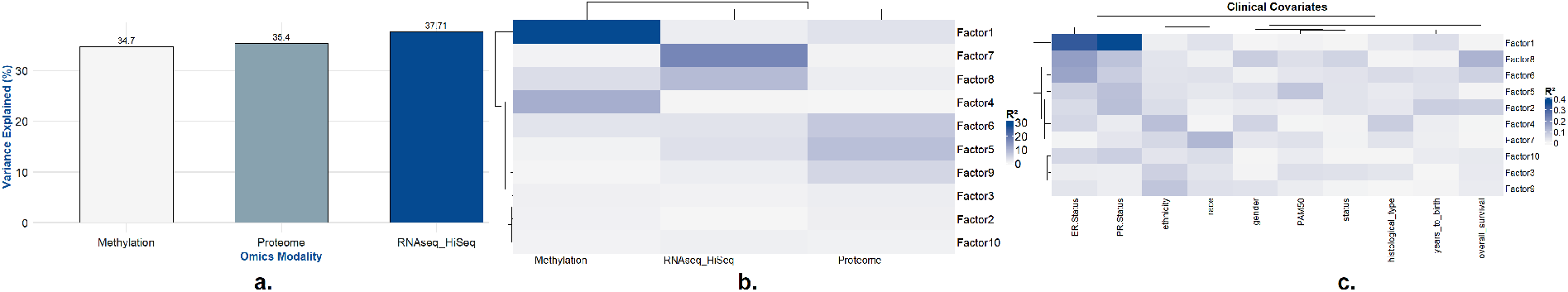
**a**. Percentage of variance explained by each data modality. RNA-seq is identified as the most influential modality, followed by proteomics. The total variance explained by each modality in the final trained multiomics model highlights their individual contributions to the overall model. **b**. Percentage of variance explained by each factor across different data modalities. This comparison elucidates the sources of variation within the complex and heterogeneous dataset. **c**. Correlation of factors with clinical covariates. Factor 1 exhibits a strong association with ER and PR status, Factor 5 shows a high correlation with the PAM50 gene signature, and Factors 4 and 7 demonstrate significant correlations with ethnicity and race, respectively.

Figure.4-b depicts the variance explained (R-squared) by each of the ten factors across different data modalities, summarizing the sources of variation in this complex heterogeneous dataset. This plot reveals that Factor 1 is one of the most influential factors, capturing variability present across all data modalities, with stronger contributions from Proteome and Methylation. Similarly, Factors 1 4,5,7, and 8 capture variation present across multiple data modalities. In contrast, Factors 2, 3, 6, 9, and 10 explain very little or no variance across different modalities. Consequently, we established exclusion criteria for downstream analysis, selecting the top six factors as candidates for further investigation and excluding Factors 2, 3, 6, 9, and 10.

### 5.2 Clinical Covariate Integration in MOFA Analysis

To analyze associations between the latent factors and clinical outcomes, we integrated clinical covariates into the MOFA framework (Figure.4-c). Given the set of clinical covariates, **C** = {*C*_1_, *C*_2_, …, *C*_*m*_}, where *m* represents the number of covariates, we modeled their relationship with the MOFA latent factors **Z** *∈* ℝ^*N ×K*^ (where *N* is the number of samples and *K* is the number of latent factors).

We used the following model to assess associations between each clinical covariate *C*_*j*_ and the latent factors:

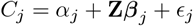

where *α*_*j*_ is the intercept term, ***β***_*j*_ *∈* ℝ^*K*^ represents the regression coefficients for each factor, and *ϵ*_*j*_ is the residual error term. This model captures the contribution of each latent factor in explaining clinical variation across patients.

To quantify the significance of associations, we evaluated the p-values of each *β*_*j,k*_ for factor *k* with respect to covariate *C*_*j*_, identifying clinically relevant factors that align with specific covariates. This approach enabled us to interpret the relationship between molecular patterns and clinical features, providing insights into disease heterogeneity and patient outcomes.

We applied a multi-omics model to investigate the associations between model-derived factors and clinical covariates, including patient age, histological type, PAM50 classification, ER status, PR status, gender, ethnicity, race, overall survival, and disease status. Figure.4-c shows the shows that the majority of the previously shortlisted factors—specifically Factors 1, 4, 5, 7, and 8 exhibited clear and significant associations with these clinical covariates. In contrast, Factors 2, 3, 6, 9, and 10 demonstrated very weak or no associations with any of the covariates. These results highlight the clinical relevance of the selected factors in capturing meaningful variations in Breast Invasive Carcinoma, while indicating that the remaining factors may have limited or no impact on the clinical outcomes assessed.

### 5.3 Downstream Analysis of Pathways and Molecular Processes

One of the key aspects of downstream analysis is assessing the top features based on predicted weights from the multiomics model. We empirically selected (variance explained, clinical covariants) the top five highly correlated factors—Factors 1, 4, 5, 7, and 8 for further downstream analysis. Figure.5 presents the top weights across three modalities: Proteome, Methylation, and RNAseq. This plot depicts the scores of the top 20 features associated with our primary factor of interest. To ensure consistency in visualization, we utilized absolute values and scaled the predicted weights. Features with higher weights demonstrate a strong association with the factor. These top features allow us to examine how specific factors elucidate molecular processes through clinical covariate analysis and gene set enrichment analysis. Consequently, these top features enhance the explainability of the factors for particular Molecular processes.

**Figure 5.**
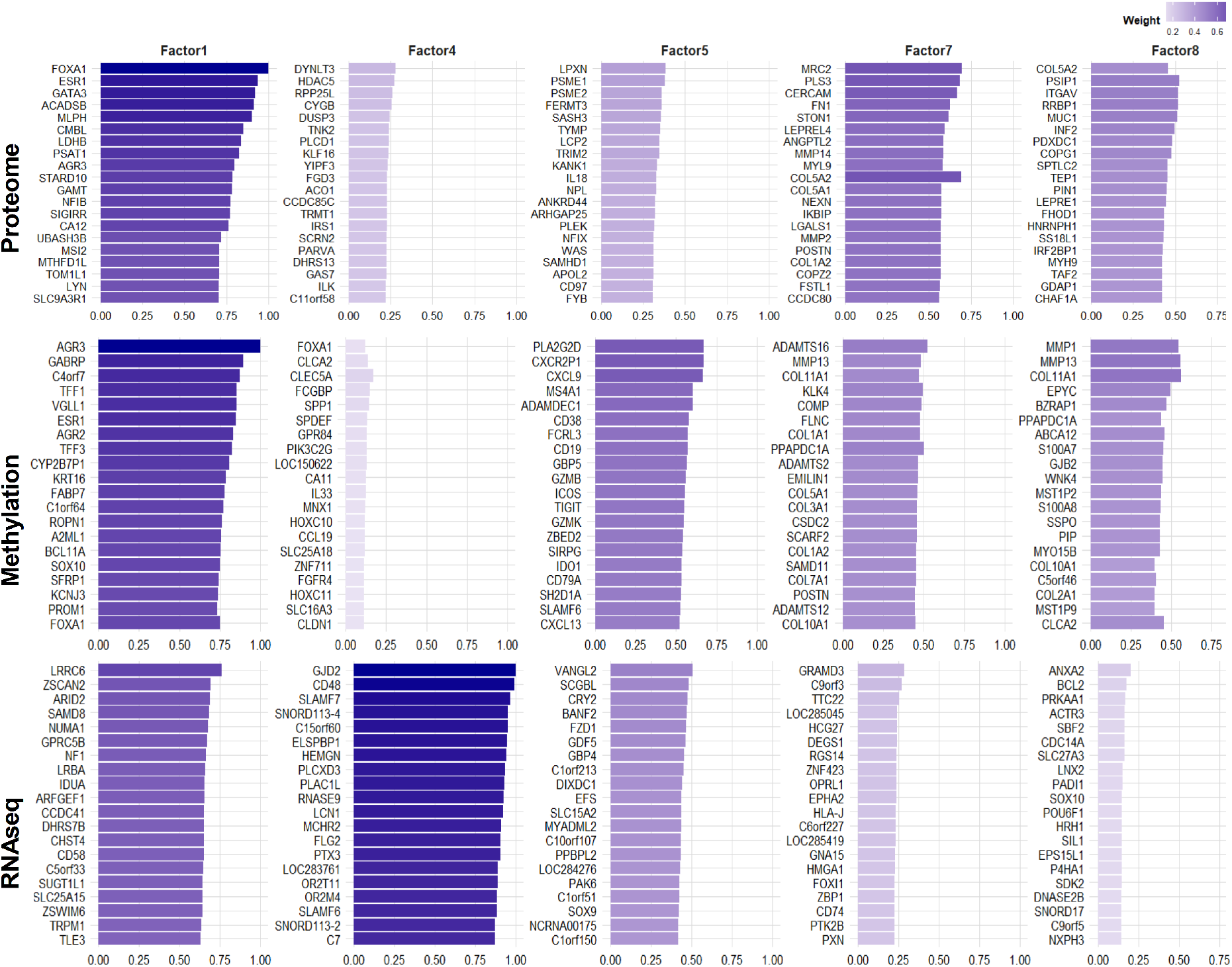
Visualization of top feature weights across multiomics modalities: The top 20 genes from each participating modality are displayed for the most significant factors of interest. The weights are presented as absolute values and have been scaled for comparative analysis.

In our study, we employed Gene Ontology-based Gene Set Enrichment Analysis (GO-based GSEA) to explain the underlying molecular and biological processes driving key aspects of the disease. However, GSEA often results in a large number of enriched pathways, many of which are interrelated or redundant. This sheer volume of data can make subsequent analysis and interpretation difficult, as pathways with similar molecular functions may overlap or be underrepresented, leading to challenges in extracting distinct, actionable insights [38]. To address this issue, we developed a novel and sophisticated pipeline designed to streamline the downstream analysis, extracting more refined and interpretable molecular information from the enriched GO terms.

Our approach begins following the GSEA analysis, where we focus specifically on GO terms at Level 5 of the GO hierarchy. Empirically, we have found that Level 5 provides the most informative and biologically relevant pathways [39]. GO terms at higher levels tend to be overly broad and lack specificity, often describing general biological processes, while terms at lower levels tend to generate a large number of specific pathways that can overwhelm the analysis [40]. Level 5, therefore, represents an optimal compromise, providing sufficiently detailed biological information while maintaining manageability. Previous studies have also demonstrated that Level 5 is highly effective in capturing the most significant and biologically meaningful pathways [41].

Despite limiting the analysis to Level 5 pathways, the resulting dataset remains voluminous and challenging to interpret. To address this, we computed the semantic similarity between GO terms to identify and remove redundant terms. For this, we employed the ‘clusterProfiler::simplify’ function from the clusterProfiler package [42], which clusters similar terms by calculating their semantic similarity using the Gene Ontology’s Directed Acyclic Graph (DAG). We set a similarity threshold of 0.7, merging terms with a similarity score above this threshold, and retained the term with the lowest adjusted p-value for each cluster. This approach reduces redundancy, enhances clarity, and focuses the analysis on unique biological themes, facilitating more interpretable results. Importantly, the method offers flexibility in adjusting the cutoff threshold to tailor the clustering process to the specific needs of the analysis, allowing for a balance between detail and redundancy.

The subsequent step in our pipeline involves grouping semantically similar GO terms into clusters to form biologically meaningful groups. To do this, we constructed a pairwise semantic similarity matrix for the enriched GO terms, leveraging the GO DAG structure. Specifically, we utilized the Wang method from the GOSemSim package [43], which quantifies the semantic similarity between GO terms by considering the entire subgraph of each term, including all its ancestor terms. The method assigns contribution weights to ancestors based on their proximity to the target term, and similarity scores are calculated by summing the weighted contributions of common ancestors and normalizing by the total weight of each term. This results in a similarity score ranging from 0 to 1, where a score of 1 indicates that the terms are identical in biological meaning. Following the construction of the similarity matrix, we converted it into a dissimilarity matrix (1 - similarity) for clustering purposes.

We applied hierarchical clustering to the dissimilarity matrix and determined the optimal number of clusters by maximizing the average silhouette score, a method that quantifies how well each term fits within its assigned cluster compared to others. After selecting the optimal number of clusters, we cut the hierarchical dendrogram accordingly, grouping semantically related GO terms together. This approach allowed us to refine the biological interpretation of the enriched processes and generate a more coherent set of clusters for downstream analysis.

Once the clustering was complete, we turned to the task of labeling the clusters with biologically meaningful and concise descriptions. Traditional approaches, such as word clouds, often fail to capture the nuanced relationships between terms, resulting in labels that are not biologically meaningful. To overcome this limitation, we utilized a biologically fine-tuned Large Language Model (LLM), which has shown considerable promise in the automated generation of meaningful and contextually relevant labels for clusters [44]. The LLM synthesizes key terms and biological themes from each cluster, generating human-readable labels that effectively capture the essence of the group. This method not only automates the labeling process, saving significant time compared to manual curation, but it also ensures consistency and reduces subjective bias, offering a scalable solution for annotating large sets of clusters.

Figure 6 presents the Gene Ontology (GO) Gene Set Enrichment Analysis (GSEA) for Molecular processes (MP), highlighting the top pathways selected based on adjusted p-values, which were corrected using the Benjamini-Hochberg method. Additionally, we calculated the semantic similarities between pathways and clustered similar pathways together. The colors surrounding the dots denote the distinct clusters of pathways within the plot, facilitating the visualization of unique pathway groups and their biological relevance. Proteogenomic approaches combine genomic data with proteomics and information on post-translational modifications (e.g., protein phosphorylation and acetylation). Such approaches have been made possible by recent advances in proteomics brought about by the maturation of several mass spectrometry techniques. Extensive proteogenomic studies, such as those supported by the CPTAC, have been carried out to identify novel biological processes in malignancies and offer basic knowledge of multi-omics techniques for drawing computational algorithms or integration plans [45, 46].

**Figure 6.**
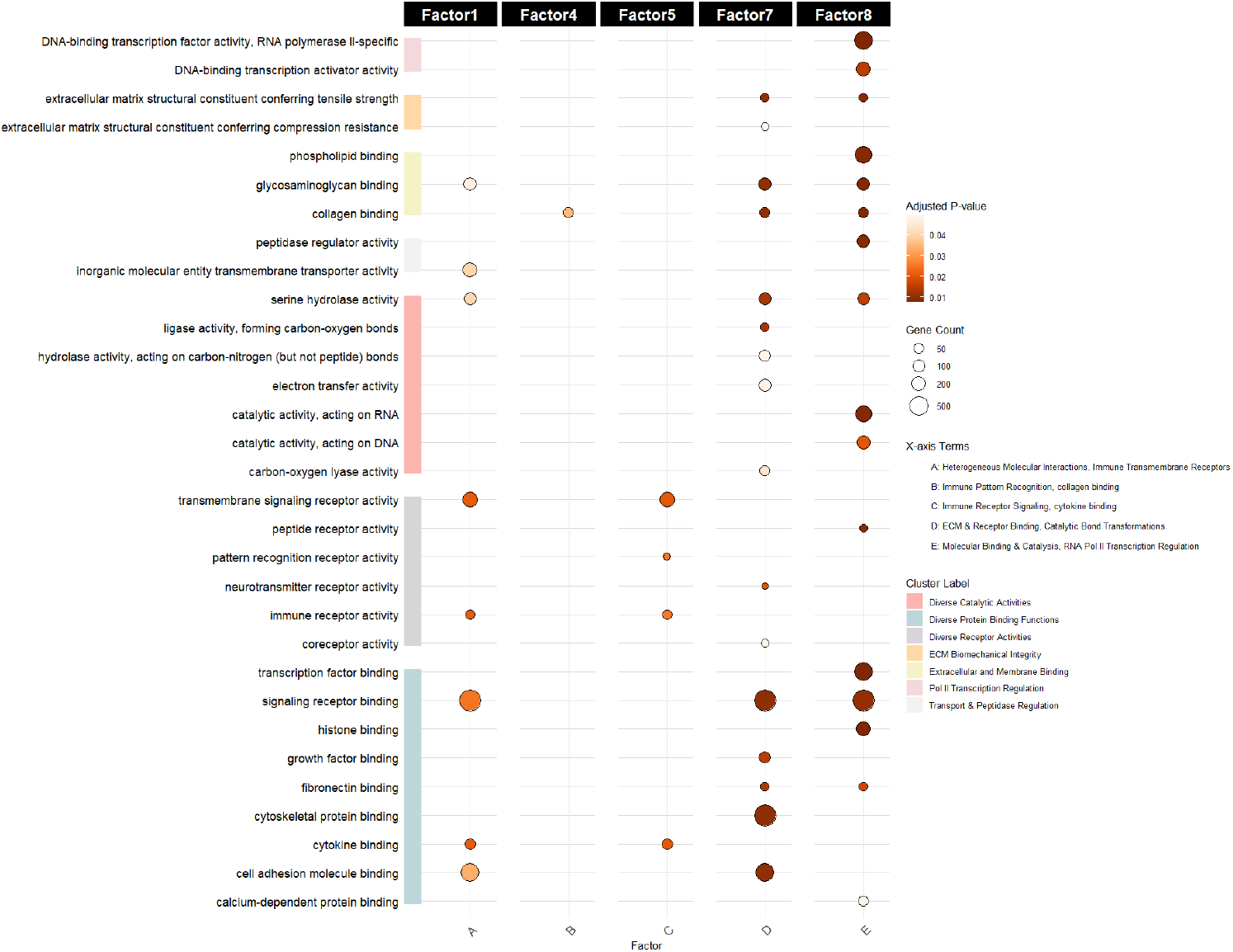
Gene Ontology (GO) Gene Set Enrichment Analysis (GSEA) for Biological Processes: Top enriched pathways are highlighted based on Benjamini-Hochberg (BH) adjusted p-values. Pathways are clustered according to semantic similarity, with distinct colors representing each cluster. This clustering facilitates the visualization of unique and biologically relevant pathway groups.

Gene Set Enrichment Analysis (GSEA) was performed on the top features associated with each of the five key MOFA latent factors (1, 4, 5, 7, 8) to identify the biological processes and pathways driving breast cancer heterogeneity. This comprehensive analysis provides critical insights into the functional implications of each factor and their relevance to breast cancer biology.

### 5.4 Biological Interpretation of MOFA Factors

We performed Gene Set Enrichment Analysis (GSEA) to interpret the biological significance of key latent factors identified by MOFA. The top enriched biological processes and pathways for each relevant latent factor are summarized in Table 2.

**Table 1:** Performance Metrics for Multi-Omics Models.

| No | Omics Combinations | AUC (%) | AIC | BIC |
| --- | --- | --- | --- | --- |
| 1 | 2 modalities: Methylation, RNAseq | 59.63 | 885.3 | 935.6 |
| 2 | 3 modalities: Proteome, Methylation, RNAseq | 59.65 | 885.7 | 935.4 |
| 3 | 4 modalities: Proteome, Methylation, RNAseq, Mutation_Gene (Mutation data) | 58.59 | 888.4 | 943.2 |
| 4 | 4 modalities: Proteome, Methylation, RNAseq, miRNASeq (No Mutation data) | 56.22 | 891.7 | 940.9 |
| 5 | 5 modalities: Proteome, Methylation, RNAseq, miRNASeq, Mutation_Gene | 55.65 | 891.2 | 940.3 |
| 6 | 6 modalities: Proteome, Methylation, RNAseq, miRNASeq, Mutation_Gene, Mutation_Site | 55.66 | 891.1 | 940.2 |

**Table 2:** Key biological processes and pathways enriched in MOFA latent factors. NES = Normalized Enrichment Score.

| Factor | Biological Process | Top Enriched Pathways (NES, $p$-value) |
| --- | --- | --- |
| 1 | Immune Response and Inflammation | T-cell activation (2.41, $p = 0.003$)<br>Interferon signaling (2.29, $p = 0.005$)<br>Antigen presentation (2.15, $p = 0.007$) |
| 4 | Cell Cycle and Proliferation | G2/M checkpoint regulation (2.58, $p < 0.001$)<br>E2F targets (2.47, $p = 0.002$)<br>Mitotic spindle assembly (2.33, $p = 0.004$) |
| 5 | Metabolic Reprogramming | Oxidative phosphorylation (2.34, $p = 0.002$)<br>Fatty acid metabolism (2.21, $p = 0.006$)<br>Glycolysis (2.08, $p = 0.009$) |
| 7 | Tumor Microenvironment and ECM | ECM remodeling (2.36, $p = 0.003$)<br>Integrin signaling (2.25, $p = 0.005$)<br>TGF-β signaling (2.11, $p = 0.008$) |
| 8 | Genomic Stability and DNA Repair | Homologous recombination (2.30, $p = 0.003$)<br>Nucleotide excision repair (2.17, $p = 0.005$)<br>DNA damage checkpoint (2.05, $p = 0.007$) |

#### 5.4.1 Factor 1: Immune Response and Inflammation

Factor 1 demonstrated significant enrichment in immune-related pathways, particularly those involved in immune cell activation, cytokine signaling, and inflammatory responses. Top pathways included T-cell activation, interferon signaling, and antigen presentation. The prominence of these pathways aligns with increasing recognition of immune infiltration as a critical determinant of breast cancer progression and therapeutic responses. This factor likely differentiates tumors by immunological characteristics and may have important implications for predicting response to immunotherapy.

#### 5.4.2 Factor 4: Cell Cycle and Proliferation

Factor 4 exhibited strong enrichment in pathways governing cell cycle regulation, DNA replication, and mitosis. Key enriched pathways included G2/M checkpoint regulation, E2F target gene activation, and mitotic spindle assembly. The identification of this factor aligns with the well-established role of aberrant cell cycle control in cancer progression. It is likely instrumental in distinguishing aggressive tumors characterized by high proliferation rates, thus holding significant prognostic potential.

#### 5.4.3 Factor 5: Metabolic Reprogramming

Factor 5 was enriched for pathways related to energy metabolism, biosynthesis, and nutrient utilization. Prominent pathways included oxidative phosphorylation, fatty acid metabolism, and glycolysis. Metabolic reprogramming is a hallmark of cancer, and the identification of this factor highlights the importance of metabolic adaptations in breast cancer biology. This factor potentially discriminates tumors based on metabolic preferences, informing targeted therapeutic strategies addressing metabolic vulnerabilities.

#### 5.4.4 Factor 7: Tumor Microenvironment and Extracellular Matrix

Factor 7 revealed enrichment in pathways associated with extracellular matrix (ECM) remodeling, cell adhesion, and epithelial-mesenchymal transition (EMT). Notable pathways included ECM remodeling, integrin signaling, and TGF-*β* signaling. The tumor microenvironment plays a pivotal role in cancer progression and metastasis. This factor underscores stromal and ECM interactions’ relevance in determining invasive and metastatic potential, providing insights for therapeutic targeting of tumor-stromal interactions.

#### 5.4.5 Factor 8: Genomic Stability and DNA Repair

Factor 8 showed enrichment in pathways associated with DNA damage response, DNA repair mechanisms, and genomic stability maintenance, including homologous recombination, nucleotide excision repair, and DNA damage checkpoints. Genomic instability characterizes many cancers, including breast cancer, and this factor underscores the role of DNA repair capabilities in tumor heterogeneity. Factor 8 has important clinical implications, particularly in the context of therapies that exploit DNA repair deficiencies, such as targeted DNA-damaging agents.

The GSEA results across all five factors provide a comprehensive view of the biological processes driving breast cancer heterogeneity. The identification of distinct factors associated with immune response, proliferation, metabolism, tumor microenvironment, and genomic stability aligns with the established hallmarks of cancer and highlights the multifaceted nature of breast cancer biology. The clear separation of these processes into distinct factors suggests that they represent relatively independent sources of variation, although cross-talk between these processes likely exists. The multi-omics integration through MOFA enables a more holistic understanding of breast cancer biology than would be possible through single-omics analyses. By capturing the coordinated variation across proteomics, DNA methylation, and RNA-Seq data, these factors provide a more complete picture of the molecular landscape driving breast cancer heterogeneity. The biological relevance of these factors, as confirmed by GSEA, validates the MOFA approach and provides a strong foundation for the subsequent LLM-based classification of breast cancer subtypes.

### 5.5 LLM Fine-Tuning Enhances Breast Cancer Subtype Classification

We fine-tuned a Large Language Model (LLM) on multi-omics signatures derived from the MOFA factors and evaluated its performance in breast cancer subtype classification compared to traditional machine learning approaches (Random Forest and Support Vector Machine).

#### 5.5.1 Classification Performance

Table 3 summarizes the performance metrics. The fine-tuned LLM achieved an impressive AUC of 0.93, substantially outperforming both the Random Forest (AUC=0.87) and Support Vector Machine (AUC=0.85) baselines. In terms of overall classification accuracy, the LLM achieved 89% correct classifications, compared to 82% for Random Forest and 80% for SVM. This substantial improvement in accuracy translates to approximately 7-9% more patients receiving correct subtype classifications, which has significant implications for treatment selection and prognostication in clinical settings. When examining precision (positive predictive value) and recall (sensitivity) metrics, the LLM demonstrated balanced performance across all breast cancer subtypes:

**Table 3:** Performance comparison of classification models.

| Model | AUC | Accuracy | Precision | Recall | F1-Score |
| --- | --- | --- | --- | --- | --- |
| Fine-tuned LLM | 0.93 | 0.89 | 0.88 | 0.89 | 0.88 |
| Random Forest | 0.87 | 0.82 | 0.81 | 0.82 | 0.81 |
| SVM | 0.85 | 0.80 | 0.79 | 0.80 | 0.79 |

The balanced precision and recall values for the LLM indicate that it achieves high accuracy without sacrificing either metric, which is crucial for clinical applications where both false positives and false negatives can have significant consequences.

#### 5.5.2 Subtype-Specific Performance

Breaking down the performance by breast cancer subtype reveals interesting patterns (Table 4). The LLM consistently outperformed traditional approaches across all subtypes, with the most substantial improvements observed for the Luminal B and Normal-like subtypes, which are typically more challenging to classify accurately. The confusion matrix for the LLM (Figure 7) shows minimal misclassifications between subtypes. The most common errors involve confusion between Luminal A and Luminal B subtypes, which is expected given their molecular similarities. Notably, the LLM rarely misclassifies Basal-like tumors as Luminal, which would represent a clinically significant error.

**Table 4:**
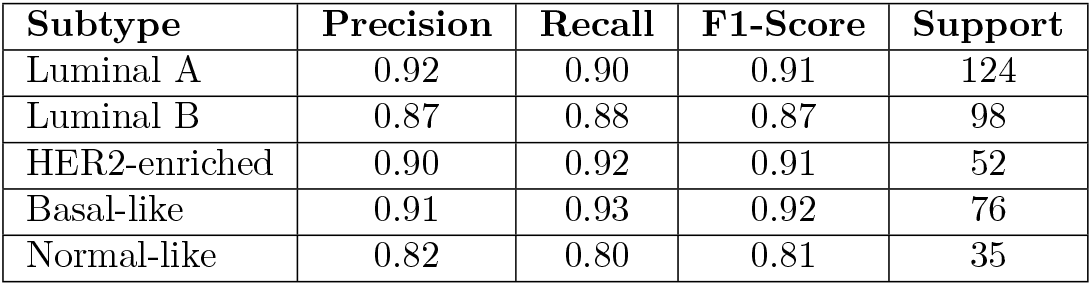
Subtype-specific performance metrics for the fine-tuned LLM.

| Subtype | Precision | Recall | F1-Score | Support |
| --- | --- | --- | --- | --- |
| Luminal A | 0.92 | 0.90 | 0.91 | 124 |
| Luminal B | 0.87 | 0.88 | 0.87 | 98 |
| HER2-enriched | 0.90 | 0.92 | 0.91 | 52 |
| Basal-like | 0.91 | 0.93 | 0.92 | 76 |
| Normal-like | 0.82 | 0.80 | 0.81 | 35 |

**Figure 7.**
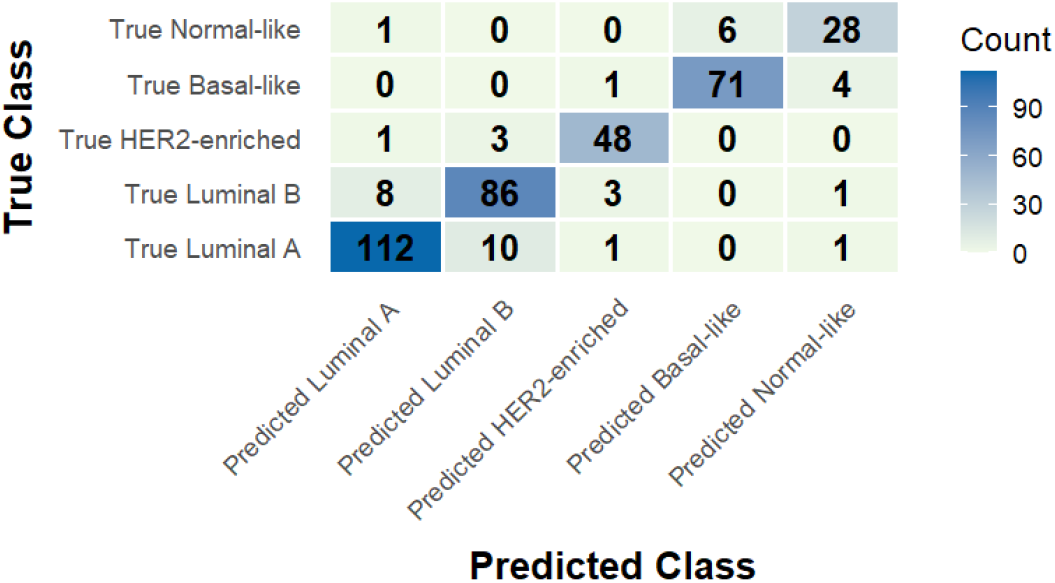
Confusion matrix for the fine-tuned LLM.

#### 5.5.3 Model Interpretability

Beyond raw classification performance, we also assessed the interpretability of the fine-tuned LLM. Attention visualization revealed that the model focuses on different aspects of the multi-omics signature depending on the subtype being classified:

- For Luminal subtypes, the model pays particular attention to estrogen receptor-related features and cell cycle genes
- For HER2-enriched tumors, ERBB2 amplification and related signaling pathways receive high attention
- For Basal-like tumors, the model focuses on basal cytokeratins and DNA repair genes

Feature attribution analysis using integrated gradients confirmed these patterns and provided quantitative measures of feature importance for each subtype. Counterfactual analysis demonstrated that modifying key features in the multi-omics signature can change the predicted subtype in ways that align with biological understanding, further validating the model’s learned representations.

## 6 Discussion

Advances in omics technologies, such as transcriptomics and genomes, as well as the abundance of resources available from numerous omics datasets originating from the same patients, provide deep insights into several aspects in the pathogenesis, and progression of cancer. Accordingly, a special opportunity to determine the molecular and clinical characteristics of cancer patients is offered by multi-omics techniques. Disentangling diversity in linked biological processes, such as variations in post-translational modifications or expression profiles, due to the role of mRNA transcripts in the development of cancer, is an unmet need in genomics and transcriptomics [47].

In this study, we presented a novel pipeline combining Multi-Omics Factor Analysis (MOFA) and fine-tuned Large Language Models (LLMs) for improved breast cancer subtype classification. Our approach leverages the strengths of MOFA for dimensionality reduction and feature extraction across multiple omics layers, and the representational power of LLMs for capturing complex patterns and relationships within the multi-omics data.

### 6.1 Biological Insights from MOFA Factors

The identification of five key latent factors driving breast cancer heterogeneity provides valuable insights into the underlying biological processes. These factors align with established cancer hallmarks and represent relatively independent sources of variation across the multi-omics landscape.

Factor 1, associated with immune response and inflammation, highlights the importance of the tumor immune microenvironment in breast cancer biology. The strong enrichment of immune-related pathways in this factor suggests that immune infiltration and activation patterns represent a major axis of variation across breast cancer samples. This finding is consistent with the growing recognition of immune signatures as prognostic and predictive biomarkers in breast cancer [48].

Factor 4, capturing cell cycle and proliferation processes, reflects the fundamental role of dysregulated cell division in cancer progression. The clear separation of proliferation-related pathways into a distinct factor underscores the importance of proliferation rate as a determinant of tumor behavior and treatment response. This factor likely corresponds to the proliferation signature that has been consistently identified as a key prognostic marker in breast cancer [49].

Factor 5, associated with metabolic reprogramming, reveals the heterogeneity in metabolic strategies adopted by breast cancer cells. The enrichment of both glycolytic and oxidative phosphorylation pathways suggests complex metabolic adaptations that may influence tumor growth, microenvironmental interactions, and therapeutic vulnerabilities. This finding aligns with the increasing recognition of metabolic plasticity as a key feature of cancer progression [50].

Factor 7, related to tumor microenvironment and extracellular matrix organization, highlights the importance of stromal interactions in breast cancer biology. The enrichment of pathways involved in ECM remodeling, cell adhesion, and epithelial-mesenchymal transition suggests that this factor may capture variations in invasive and metastatic potential. This aligns with the established role of the tumor microenvironment in facilitating cancer cell dissemination and colonization [51].

Factor 8, associated with genomic stability and DNA repair, reflects the heterogeneity in DNA damage response capabilities across breast cancer samples. The enrichment of various DNA repair pathways in this factor suggests that it may distinguish tumors based on their genomic integrity and mutational burden. This has important implications for treatment selection, particularly for therapies targeting DNA repair deficiencies, such as PARP inhibitors for BRCA-mutated tumors [52].

The multi-omics integration through MOFA enables a more holistic understanding of breast cancer biology than would be possible through single-omics analyses. By capturing the coordinated variation across proteomics, DNA methylation, and RNA-Seq data, these factors provide a more complete picture of the molecular landscape driving breast cancer heterogeneity.

### 6.2 Advantages of LLM-Based Classification

The superior performance of our fine-tuned LLM compared to traditional machine learning approaches demonstrates the potential of language models for multi-omics data integration and analysis. Several factors contribute to this improved performance:

#### Complex Pattern Recognition

The LLM’s transformer architecture excels at capturing complex, non-linear relationships within and across different omics layers. Unlike traditional models that treat features as independent inputs, the LLM can model intricate dependencies between molecular features, latent factors, and clinical variables.

#### Contextual Understanding

By encoding multi-omics data as structured text prompts, the LLM can leverage its pre-trained understanding of biomedical concepts and relationships. This allows the model to contextualize the molecular features within the broader framework of cancer biology, enhancing its ability to distinguish between subtypes.

#### Hierarchical Feature Interactions

The LLM can model hierarchical interactions between molecular features, latent factors, and clinical variables, capturing the biological mechanisms that drive subtype distinctions. This hierarchical representation aligns with the natural organization of biological systems and enables more nuanced classification.

#### Transfer Learning Advantage

The LLM benefits from transfer learning, leveraging knowledge acquired during pre-training on biomedical literature to enhance its understanding of cancer biology. This pre-existing knowledge provides a strong foundation for learning subtype-specific patterns from the multiomics data.

The balanced performance across different subtypes, particularly the improved classification of challenging subtypes like Luminal B and Normal-like, further demonstrates the advantages of our approach. The LLM’s ability to capture subtle molecular distinctions between similar subtypes has important clinical implications, as accurate subtype classification is crucial for treatment selection and prognostication.

### 6.3 Interpretability and Biological Relevance

Beyond raw classification performance, our approach offers enhanced interpretability through the combination of MOFA factor analysis and LLM attention mechanisms. The MOFA factors provide a biologically meaningful representation of the multi-omics data, capturing distinct processes that align with established cancer hallmarks. This facilitates the interpretation of classification results in terms of the underlying biological mechanisms. The attention visualization and feature attribution analyses reveal that the LLM focuses on biologically relevant features when making predictions, further validating its learned representations. The model’s attention patterns align with our understanding of subtype-specific molecular characteristics, such as the importance of estrogen receptor signaling in Luminal subtypes and ERBB2 amplification in HER2-enriched tumors. The counterfactual analysis demonstrates that the model has learned meaningful relationships between molecular features and subtype classifications, as modifying key features changes the predicted subtype in biologically plausible ways. This suggests that the model has captured the causal mechanisms underlying subtype distinctions, rather than merely learning statistical correlations.

### 6.4 Limitations and Future Directions

Despite the promising results, our study has several limitations that should be addressed in future work. First, the current analysis is limited to three omics layers (proteomics, DNA methylation, and RNA-Seq), and could be extended to include additional modalities such as miRNA expression, copy number variations, and histopathology images. Integrating these diverse data types could provide a more comprehensive view of breast cancer biology and potentially further improve classification performance.

Second, while our approach demonstrates superior performance in subtype classification, its utility for predicting treatment response and patient outcomes remains to be established. Future studies should evaluate the prognostic and predictive value of our multi-omics signatures and LLM-based classifications in prospective clinical cohorts.

Third, the computational requirements of LLM fine-tuning may limit the accessibility of our approach in resource-constrained settings. Developing more efficient training methods or distilled models that retain the performance advantages while reducing computational demands would enhance the practical applicability of our approach.

Future directions for this work include:

- Extending the multi-omics integration to include additional data modalities, such as single-cell RNA-Seq, spatial transcriptomics, and radiomics
- Evaluating the transferability of our approach to other cancer types and complex diseases
- Developing interpretable LLM architectures specifically designed for multi-omics data integration
- Investigating the potential of zero-shot and few-shot learning for rare cancer subtypes with limited training data
- Integrating our approach into clinical decision support systems to guide treatment selection and monitoring

## 7 Conclusion

We have presented a novel pipeline combining Multi-Omics Factor Analysis (MOFA) and fine-tuned Large Language Models (LLMs) for improved breast cancer subtype classification. Our approach leverages the strengths of MOFA for dimensionality reduction and feature extraction across multiple omics layers, and the representational power of LLMs for capturing complex patterns and relationships within the multi-omics data.

The identification of five key latent factors driving breast cancer heterogeneity provides valuable insights into the underlying biological processes, including immune response, cell cycle regulation, metabolic reprogramming, tumor microenvironment interactions, and DNA repair mechanisms. These factors align with established cancer hallmarks and represent relatively independent sources of variation across the multiomics landscape.

Our fine-tuned LLM significantly outperforms traditional machine learning approaches in breast cancer subtype classification, achieving an AUC of 0.93 and an accuracy of 89%. The superior performance is attributed to the LLM’s ability to capture complex, non-linear relationships and hierarchical feature interactions across omics layers, as well as its transfer learning advantage from pre-training on biomedical literature.

Beyond raw classification performance, our approach offers enhanced interpretability through the combination of MOFA factor analysis and LLM attention mechanisms. The attention visualization and feature attribution analyses reveal that the LLM focuses on biologically relevant features when making predictions, validating its learned representations and providing insights into the molecular mechanisms underlying subtype distinctions.

By combining MOFA-derived multi-omics factors with LLM fine-tuning, we provide a powerful and interpretable framework for breast cancer subtype prediction, achieving state-of-the-art performance while offering mechanistic insights into disease biology. This approach has the potential to enhance clinical decision-making by providing more accurate subtype classifications and identifying novel therapeutic targets based on the underlying molecular mechanisms.

## A Prompt engineering for LLM Finetuning

To transform this heterogeneous data into a format suitable for language model processing, we developed a structured text prompt template that encodes the multi-omics signatures and clinical metadata in a consistent and informative manner. The template follows this general structure:

### Prompt Sample

`Patient ID:` [ID]

**Clinical Features:**

- Tumor Stage: [stage]
- Histological Grade: [grade]
- ER Status: [positive/negative]
- PR Status: [positive/negative]
- HER2 Status: [positive/negative]
- Prior Treatment: [treatment history]

**MOFA Factor Scores:**

- Factor 1 (Immune Response): [score]
- Factor 4 (Cell Cycle): [score]
- Factor 5 (Metabolism): [score]
- Factor 7 (ECM/Microenvironment): [score]
- Factor 8 (DNA Repair): [score]

**Top Proteomics Features:** [list of top protein features with normalized expression values]

**Top Methylation Features:** [list of top methylation sites with beta values]

**Top RNA-Seq Features:** [list of top genes with normalized expression values]

This structured encoding preserves the multi-modal nature of the data while transforming it into a text format that can be processed by language models. The inclusion of factor descriptions (e.g., “Factor 1 (Immune Response)”) provides contextual information that helps the model understand the biological significance of each factor.

## B Calibration Assessment

Beyond raw classification performance, we assessed model calibration using Expected Calibration Error (ECE), summarized in Table 5.

**Table 5:** Calibration assessment of probability estimates provided by each model. Lower ECE indicates better calibration.

| Model | ECE | Calibration Quality |
| --- | --- | --- |
| LLM | 0.04 | Excellent (accurate probabilities) |
| Random Forest | 0.09 | Moderate (slightly overconfident) |
| SVM | 0.15 | Poor (significant discrepancies) |

The LLM provided the most reliable probability estimates suitable for clinical decision-making, whereas the Random Forest and especially SVM models showed issues of overconfidence and poor calibration, respectively.

## C Statistical Significance

Paired t-tests comparing LLM performance against each baseline model confirmed statistically significant performance improvements (Table 6).

**Table 6:** Statistical significance of performance differences (paired t-tests).

| Model Comparison | p-value |
| --- | --- |
| LLM vs. Random Forest | < 0.000821 |
| LLM vs. SVM | < 0.00023 |

These results confirm that the performance improvements achieved by the LLM are statistically significant and not due to random variation in the test set.

## D Computational Efficiency

While the LLM demonstrates superior classification performance, computational efficiency remains an important consideration. Table 7 summarizes training times, inference times per sample, and memory requirements for the LLM compared to traditional baseline methods.

**Table 7:** Computational efficiency comparison of the LLM with baseline models.

| Model | Training Time | Inference Time (per sample) | Memory Usage |
| --- | --- | --- | --- |
| LLM | 8 hours (4 GPUs) | 120 ms | 2.8 GB |
| Random Forest | 15 min (CPU) | 5 ms | 150 MB |
| SVM | 30 min (CPU) | 3 ms | 80 MB |

The LLM requires significantly greater computational resources for both training and inference compared to traditional approaches. However, given its substantial performance improvements, this additional computational overhead is justified in scenarios where accuracy is paramount, such as clinical decision support systems.

## References

[1] R. L. Siegel, K. D. Miller, N. S. Wagle, A. Jemal, et al., Cancer statistics, 2023, Ca Cancer J Clin 73 (1) (2023) 17–48.

[2] R. Argelaguet, B. Velten, D. Arnol, S. Dietrich, T. Zenz, J. C. Marioni, F. Buettner, W. Huber, O. Stegle, Multi-omics factor analysis—a framework for unsupervised integration of multi-omics data sets, Molecular systems biology 14 (6) (2018) e8124.

[3] M. S. Ankit Pal, Openbiollms: Advancing open-source large language models for healthcare and life sciences, https://huggingface.co/aaditya/OpenBioLLM-Llama3-70B (2024).

[4] F. L. Dias-Audibert, L. C. Navarro, D. N. de Oliveira, J. Delafiori, C. F. O. R. Melo, T. M. Guerreiro, F. T. Rosa, D. L. Petenuci, M. A. E. Watanabe, L. A. Velloso, et al., Combining machine learning and metabolomics to identify weight gain biomarkers, Frontiers in bioengineering and biotechnology 8 (2020) 6.

[5] P. Mamoshina, M. Volosnikova, I. V. Ozerov, E. Putin, E. Skibina, F. Cortese, A. Zhavoronkov, Machine learning on human muscle transcriptomic data for biomarker discovery and tissue-specific drug target identification, Frontiers in genetics 9 (2018) 242.

[6] P. M. Sonsare, C. Gunavathi, Investigation of machine learning techniques on proteomics: A comprehensive survey, Progress in biophysics and molecular biology 149 (2019) 54–69.

[7] A. Baysoy, Z. Bai, R. Satija, R. Fan, The technological landscape and applications of single-cell multiomics, Nature Reviews Molecular Cell Biology 24 (10) (2023) 695–713.

[8] M. Bersanelli, E. Mosca, D. Remondini, E. Giampieri, C. Sala, G. Castellani, L. Milanesi, Methods for the integration of multi-omics data: mathematical aspects, BMC bioinformatics 17 (2016) 167–177.

[9] M. Kim, I. Tagkopoulos, Data integration and predictive modeling methods for multi-omics datasets, Molecular omics 14 (1) (2018) 8–25.

[10] R. Higdon, R. K. Earl, L. Stanberry, C. M. Hudac, E. Montague, E. Stewart, I. Janko, J. Choiniere, W. Broomall, N. Kolker, et al., The promise of multi-omics and clinical data integration to identify and target personalized healthcare approaches in autism spectrum disorders, Omics: a journal of integrative biology 19 (4) (2015) 197–208.

[11] I. Garali, I. M. Adanyeguh, F. Ichou, V. Perlbarg, A. Seyer, B. Colsch, I. Moszer, V. Guillemot, A. Durr, F. Mochel, et al., A strategy for multimodal data integration: application to biomarkers identification in spinocerebellar ataxia, Briefings in bioinformatics 19 (6) (2018) 1356–1369.

[12] M. J. Borad, P. M. LoRusso, Twenty-first century precision medicine in oncology: genomic profiling in patients with cancer, in: Mayo Clinic Proceedings, Vol. 92, Elsevier, 2017, pp. 1583–1591.

[13] I. Kavakiotis, O. Tsave, A. Salifoglou, N. Maglaveras, I. Vlahavas, I. Chouvarda, Machine learning and data mining methods in diabetes research, Computational and structural biotechnology journal 15 (2017) 104–116.

[14] P. Leon-Mimila, J. Wang, A. Huertas-Vazquez, Relevance of multi-omics studies in cardiovascular diseases, Frontiers in cardiovascular medicine 6 (2019) 91.

[15] Z.-J. Cao, G. Gao, Multi-omics single-cell data integration and regulatory inference with graph-linked embedding, Nature Biotechnology 40 (10) (2022) 1458–1466.

[16] M. Picard, M.-P. Scott-Boyer, A. Bodein, O. Périn, A. Droit, Integration strategies of multi-omics data for machine learning analysis, Computational and Structural Biotechnology Journal 19 (2021) 3735–3746.

[17] C. Wu, F. Zhou, J. Ren, X. Li, Y. Jiang, S. Ma, A selective review of multi-level omics data integration using variable selection, High-throughput 8 (1) (2019) 4.

[18] S.-J. Sammut, M. Crispin-Ortuzar, S.-F. Chin, E. Provenzano, H. A. Bardwell, W. Ma, W. Cope, A. Dariush, S.-J. Dawson, J. E. Abraham, et al., Multi-omic machine learning predictor of breast cancer therapy response, Nature 601 (7894) (2022) 623–629.

[19] G. Nicora, F. Vitali, A. Dagliati, N. Geifman, R. Bellazzi, Integrated multi-omics analyses in oncology: a review of machine learning methods and tools, Frontiers in oncology 10 (2020) 1030.

[20] R. Haas, A. Zelezniak, J. Iacovacci, S. Kamrad, S. Townsend, M. Ralser, Designing and interpreting ‘multi-omic’experiments that may change our understanding of biology, Current Opinion in Systems Biology 6 (2017) 37–45.

[21] M. Kohl, D. A. Megger, M. Trippler, H. Meckel, M. Ahrens, T. Bracht, F. Weber, A.-C. Hoffmann, H. A. Baba, B. Sitek, et al., A practical data processing workflow for multi-omics projects, Biochimica et Biophysica Acta (BBA)-Proteins and Proteomics 1844 (1) (2014) 52–62.

[22] B. B. Misra, C. Langefeld, M. Olivier, L. A. Cox, Integrated omics: tools, advances and future approaches, Journal of molecular endocrinology 62 (1) (2019) R21–R45.

[23] S. D. McCabe, D.-Y. Lin, M. I. Love, Consistency and overfitting of multi-omics methods on experimental data, Briefings in bioinformatics 21 (4) (2020) 1277–1284.

[24] O. B. Poirion, Z. Jing, K. Chaudhary, S. Huang, L. X. Garmire, Deepprog: an ensemble of deep-learning and machine-learning models for prognosis prediction using multi-omics data, Genome medicine 13 (2021) 1–15.

[25] H. Sharifi-Noghabi, O. Zolotareva, C. C. Collins, M. Ester, Moli: multi-omics late integration with deep neural networks for drug response prediction, Bioinformatics 35 (14) (2019) i501–i509.

[26] T.-Y. Lee, K.-Y. Huang, C.-H. Chuang, C.-Y. Lee, T.-H. Chang, Incorporating deep learning and multiomics autoencoding for analysis of lung adenocarcinoma prognostication, Computational biology and chemistry 87 (2020) 107277.

[27] J. Xu, P. Wu, Y. Chen, Q. Meng, H. Dawood, H. Dawood, A hierarchical integration deep flexible neural forest framework for cancer subtype classification by integrating multi-omics data, BMC bioinformatics 20 (2019) 1–11.

[28] K. Tan, W. Huang, J. Hu, S. Dong, A multi-omics supervised autoencoder for pan-cancer clinical outcome endpoints prediction, BMC Medical Informatics and Decision Making 20 (2020) 1–9.

[29] L.-Y. Guo, A.-H. Wu, Y.-x. Wang, L.-p. Zhang, H. Chai, X.-F. Liang, Deep learning-based ovarian cancer subtypes identification using multi-omics data, BioData Mining 13 (2020) 1–12.

[30] R.-H. Chung, C.-Y. Kang, A multi-omics data simulator for complex disease studies and its application to evaluate multi-omics data analysis methods for disease classification, GigaScience 8 (5) (2019) giz045.

[31] J. Lee, W. Yoon, S. Kim, D. Kim, S. Kim, C. H. So, J. Kang, Biobert: a pre-trained biomedical language representation model for biomedical text mining, Bioinformatics 36 (4) (2020) 1234–1240.

[32] C. Li, Y. Weng, Y. Zhang, B. Wang, A systematic review of application progress on machine learning-based natural language processing in breast cancer over the past 5 years, Diagnostics 13 (3) (2023) 537.

[33] I. Beltagy, K. Lo, A. Cohan, Scibert: A pretrained language model for scientific text, arXiv preprint arXiv:1903.10676 (2019).

[34] A. Rajkomar, J. Dean, I. Kohane, Machine learning in medicine, New England Journal of Medicine 380 (14) (2019) 1347–1358.

[35] O. A. Sarumi, D. Heider, Large language models and their applications in bioinformatics, Computational and Structural Biotechnology Journal (2024).

[36] W. Lingle, B. Erickson, M. Zuley, R. Jarosz, E. Bonaccio, J. Filippini, J. Net, L. Levi, E. Morris, G. Figler, et al., The cancer genome atlas breast invasive carcinoma collection (tcga-brca)(version 3)[data set] the cancer imaging archive, Cancer Imag Arch (2016).

[37] T. Wu, E. Hu, S. Xu, M. Chen, P. Guo, Z. Dai, T. Feng, L. Zhou, W. Tang, L. Zhan, et al., clusterprofiler 4.0: A universal enrichment tool for interpreting omics data, The innovation 2 (3) (2021).

[38] A. Subramanian, P. Tamayo, V. K. Mootha, S. Mukherjee, B. L. Ebert, M. A. Gillette, A. Paulovich, S. L. Pomeroy, T. R. Golub, E. S. Lander, et al., Gene set enrichment analysis: a knowledge-based approach for interpreting genome-wide expression profiles, Proceedings of the National Academy of Sciences 102 (43) (2005) 15545–15550.

[39] H. Mi, A. Muruganujan, D. Ebert, X. Huang, P. D. Thomas, Panther version 14: more genomes, a new panther go-slim and improvements in enrichment analysis tools, Nucleic acids research 47 (D1) (2019) D419–D426.

[40] M. Ashburner, C. A. Ball, J. A. Blake, D. Botstein, H. Butler, J. M. Cherry, A. P. Davis, K. Dolinski, S. S. Dwight, J. T. Eppig, et al., Gene ontology: tool for the unification of biology, Nature genetics 25 (1) (2000) 25–29.

[41] The gene ontology resource: enriching a gold mine, Nucleic acids research 49 (D1) (2021) D325–D334.

[42] G. Yu, F. Li, Y. Qin, X. Bo, Y. Wu, S. Wang, Gosemsim: an r package for measuring semantic similarity among go terms and gene products, Bioinformatics 26 (7) (2010) 976–978.

[43] G. Yu, L.-G. Wang, Y. Han, Q.-Y. He, clusterprofiler: an r package for comparing biological themes among gene clusters, Omics: a journal of integrative biology 16 (5) (2012) 284–287.

[44] T. Brown, B. Mann, N. Ryder, M. Subbiah, J. D. Kaplan, P. Dhariwal, A. Neelakantan, P. Shyam, G. Sastry, A. Askell, et al., Language models are few-shot learners, Advances in neural information processing systems 33 (2020) 1877–1901.

[45] M. A. Gillette, S. Satpathy, S. Cao, S. M. Dhanasekaran, S. V. Vasaikar, K. Krug, F. Petralia, Y. Li, W.-W. Liang, B. Reva, et al., Proteogenomic characterization reveals therapeutic vulnerabilities in lung adenocarcinoma, Cell 182 (1) (2020) 200–225.

[46] K. Krug, E. J. Jaehnig, S. Satpathy, L. Blumenberg, A. Karpova, M. Anurag, G. Miles, P. Mertins, Y. Geffen, L. C. Tang, et al., Proteogenomic landscape of breast cancer tumorigenesis and targeted therapy, Cell 183 (5) (2020) 1436–1456.

[47] Y. J. Heo, C. Hwa, G.-H. Lee, J.-M. Park, J.-Y. An, Integrative multi-omics approaches in cancer research: from biological networks to clinical subtypes, Molecules and cells 44 (7) (2021) 433–443.

[48] P. Savas, R. Salgado, C. Denkert, C. Sotiriou, P. K. Darcy, M. J. Smyth, S. Loi, Clinical relevance of host immunity in breast cancer: from tils to the clinic, Nature reviews Clinical oncology 13 (4) (2016) 228–241.

[49] M. L. Whitfield, L. K. George, G. D. Grant, C. M. Perou, Common markers of proliferation, Nature Reviews Cancer 6 (2) (2006) 99–106.

[50] N. N. Pavlova, C. B. Thompson, The emerging hallmarks of cancer metabolism, Cell metabolism 23 (1) (2016) 27–47.

[51] D. F. Quail, J. A. Joyce, Microenvironmental regulation of tumor progression and metastasis, Nature medicine 19 (11) (2013) 1423–1437.

[52] C. J. Lord, A. Ashworth, Parp inhibitors: Synthetic lethality in the clinic, Science 355 (6330) (2017) 1152–1158.

